# Measuring and Modeling Nonlinear Interactions Between Brain Regions with fMRI

**DOI:** 10.1101/074856

**Authors:** Stefano Anzellotti, Evelina Fedorenko, Alexander J E Kell, Alfonso Caramazza, Rebecca Saxe

**Affiliations:** MIT; Harvard University

## Abstract

In the study of connectivity in large-scale networks of brain regions, a standard assumption is made that the statistical dependence between regions is univariate and linear. However, brain regions encode information in multivariate responses, and neural computations are nonlinear. Multivariate and nonlinear statistical dependence between regions is likely ubiquitous, but it is not captured by current methods. To fill this gap, we introduce a novel analysis framework: fMRI responses are characterized as points in multidimensional spaces, and nonlinear dependence is modeled using artificial neural networks. Converging evidence from multiple experiments shows that nonlinear dependence 1) models mappings between brain regions more accurately than linear dependence, explaining more variance in left-out data; 2) reveals functional subdivisions within cortical networks, and 3) is modulated by the task participants are performing.

## I. INTRODUCTION

Cognitive functions are implemented by networks of interacting brain regions. From invasive research in animal models [1], it is clear that inter-regional interactions are multivariate and nonlinear. Within each cortical region, patterns of activity across neural populations contain information, beyond what is captured by the average magnitude of response [2]. Across cortical regions, this information is transformed by nonlinear computations [3]. Thus, for example, the primate ventral visual stream can be effectively modeled by convolutional neural network models that contain layers of filters, implementing nonlinear transformations [4]. There is currently tremendous interest in capturing the structure of functional inter-regional interactions in the human brain [5, 6]. Noninvasive functional neuroimaging measures the timecourse of hemodynamic activity simultaneously across the whole brain. These timecourses, measured at rest or during task performance, can be used to identify patterns of statistical dependence between brain regions or networks [7]. Comparisons of functional connectivity patterns in hundreds or thousands of human participants have been proposed, as a means to study neurodevelopmental and psychiatric disorders. However, existing methods for capturing statistical dependence using fMRI data can only detect inter-regional dependences that are (i) univariate (i.e., based on the average magnitude of response in a region; with few exceptions [8–12], and (ii) linear. Thus, a key challenge for cognitive neuroscience is to sensitively measure multivariate and nonlinear inter-regional dependence, given the limited resolution and number of measurements of typical fMRI datasets. Here we develop and validate a measure of multivariate and nonlinear statistical dependence between human brain regions in fMRI data (Fig. 1), using artificial neural networks (‘Nonlinear Multi-variate Pattern Dependence’ or NL-MVPD). Critical to the wide applicability of this method, we validate the measure in cases with strong a priori expectations (visual object identification, auditory processing, face recognition), but also show that it is sensitive to inter-regional dependence for higher-order cognitive tasks. First, NL-MVPD can select the correct model of underlying neural computations from simulated fMRI data, when the ground truth is known. Second, NL-MVPD explains the inter-regional dependence between brain regions that were previously identified as implementing successive layers of a CNN, for auditory perception. Third, NL-MVPD recovers a plausible functional network structure for the perceptual recognition of person identity, consistent with a priori expectations. Fourth, NL-MVPD proposes a functional network structure for language processing, in the absence of strong a priori constraints. Fifth, NL-MVPD is more sensitive to changes in network interactions caused by changes in cognitive tasks than are traditional methods. Overall, we find the NL-MVPD can explain independently measured activity across brain regions better than existing methods, for both sensory and cognitive tasks. The basic approach of NL-MVPD combines the strengths of two widely (but separately) used approaches in fMRI data analysis. First, similar to multi-voxel pattern analysis (MVPA, [13]), the response of each cortical region is measured by its pattern (not its average magnitude) of activity. Using PCA, each region’s timecourse of activity is described with a set of dimensions (recurring patterns of voxel coactivations); the region’s responses are represented as the position in this multidimensional space, over time. Second, as a generalization of functional connectivity methods (14), for each pair of regions, the simultaneous statistical dependence between their multivariate timecourses is estimated, using artificial neural networks to capture nonlinear dependence. Finally, the predictive power of the learned model is tested in independent (left-out) data.

**Figure 1:**
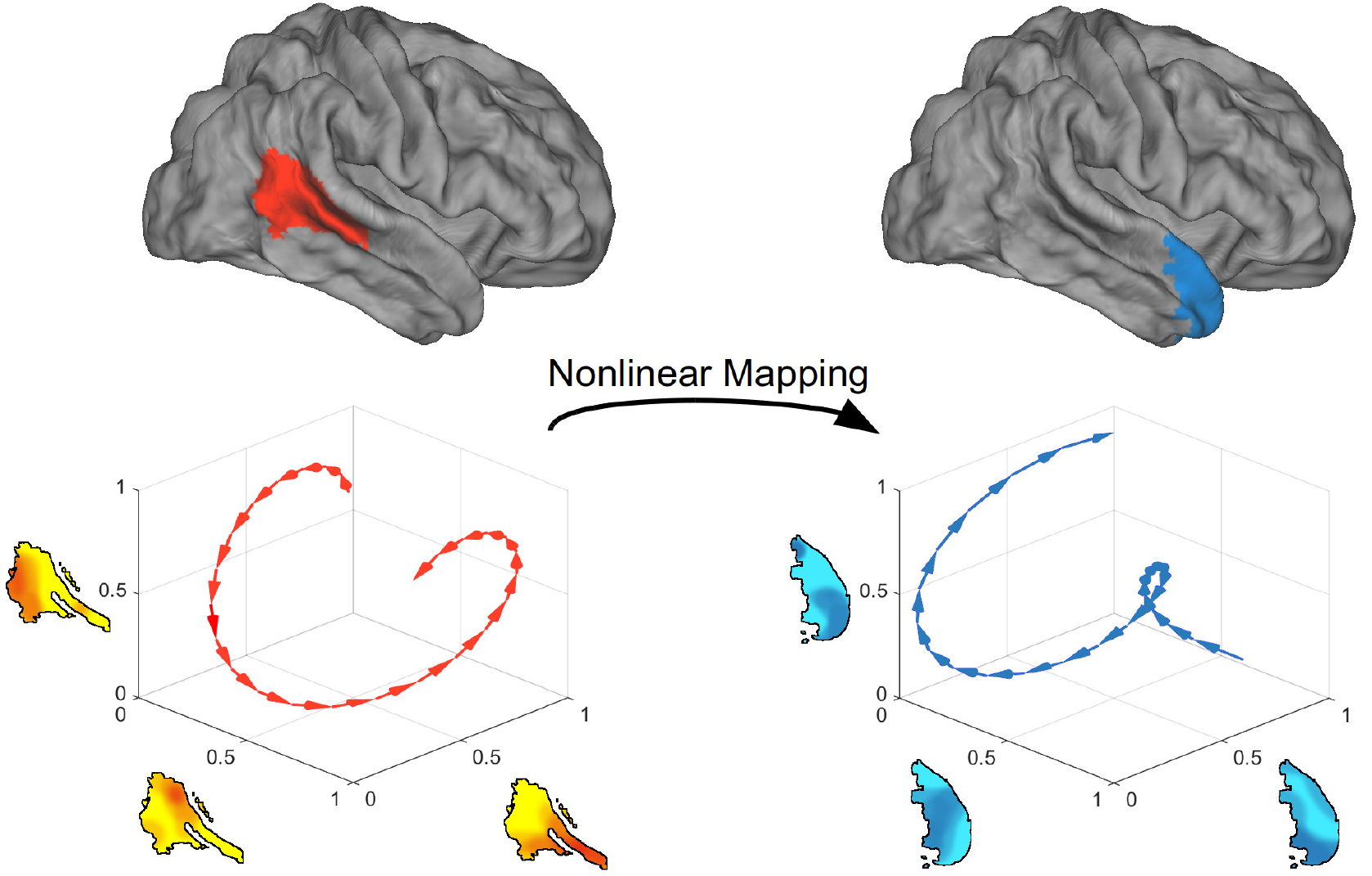
Nonlinear multivariate pattern dependence (NL-MVPD). The responses in two regions over time are described as trajectories in multidimensional spaces in which each dimension corresponds to a spatial pattern of activity. A nonlinear mapping between the two multidimensional spaces is estimated with part of the data, and its predictive power is tested in independent data.

## II. MATERIALS AND METHODS

### A. Data Simulation

The responses in the C1 and C2 layers of the HMAX model (17) were calculated for the 10 example images provided with the HMAX code (http://maxlab.neuro.georgetown.edu/hmax.html#code). Neural responses were generated assuming 2 seconds of stimulus presentation separated by 8 seconds of interstimulus interval. A simulated experiment consisted of 4 runs, in each run each image was presented twice in a randomized order. Voxel responses *V*_1_ for a first brain region *R*_1_ were generated considering each voxel as a linear combination of random proportions of each node in layer C1. The resulting values were subsequently convolved with a standard haemodynamic response function (hrf) from SPM generated using the script spm_hrf, and zero-mean gaussian noise was added. Analogously voxel responses *V*_2_ for a second brain region *R*_2_ were generated considering each voxel as a linear combination of random proportions of each node in layer C2, convolving the values with a standard hrf, and adding zero-mean gaussian noise. Each region was assumed to contain 200 voxels. Two levels of noise were used (25% and 50% of the signal’s variance), and 10 iterations of the simulation were generated for each noise level.

### B. Recovering the Ground Truth

Let’s consider brain regions *R_1_, R_2_*, in which we measured multivariate responses over time 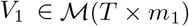 and 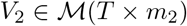, where 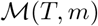 is the space of *T × m* matrices, *T* is the number of timepoints in the experiment, and *m_1_* and *m_2_* are the number of dimensions in regions *V*_1_ and *V*_2_ (which can be the number of voxels, or the number of principal components if PCA was applied). We can use the data to estimate a mapping *g: V*_1_ → *V*_2_ so that

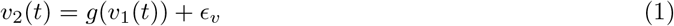

where *v_1_(t)* = *V*_1_(*t*, ·) and *v_2_(t)* = *V*_2_(*t*, ·). In this article, we estimated *g* with artificial neural networks (details in section II E) for nonlinear dependence, and with multivariate linear regression for linear dependence.

*V*_1_ and *V*_2_ are indirect measurements of the neural firing, represented by latent variables *U_1_, U_2_* in 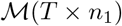 and 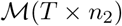 respectively, via functions *h_1_, h_2_*:

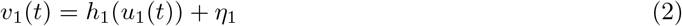

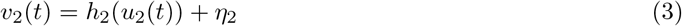

where *u_1_(t) = U_1_(t*, ·) and *u_2_(t) = U_2_(t*, ·). Furthermore, neural responses *U*_2_ are constructed from responses *U*_1_ via a function *f* - the underlying neural computation:

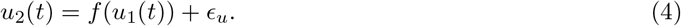

Given two computational models of the latent variables (e.g. two artificial neural networks along with a specification of which layers corresponds to which brain regions), taking the responses in the nodes of the relevant layers as 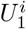 and 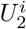 and the map between them as *f^i^* (with *i* ∈ {1, 2}), we want to determine which model provides a better account of the data.

The probability of a latent computational model 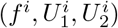 given the observable data can be described as

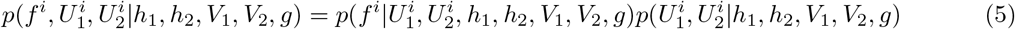

and treating 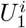 and 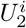 as dependent only on *V_1_, h_1_* and *V_2_, h_2_* respectively:

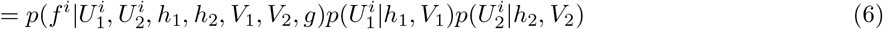

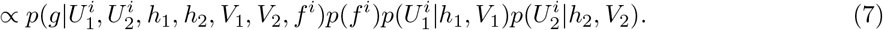

We can model *h*_1_, *h*_1_ as a composition of a linear transformation mapping neurons to voxels, and a convolution with a standard haemodynamic response function. Each neuron is considered as a ‘neuron type’ (the number of neurons in a brain region is much larger than the number of nodes in the neural network), and the weights of the linear transformation reflect the proportions of each different neuron type contained in each voxel.

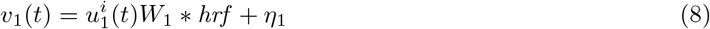

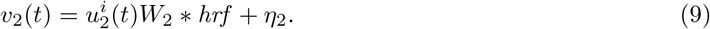

We have that

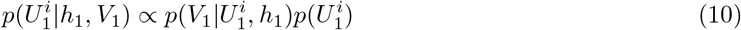

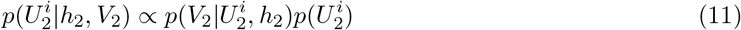

where we assume uniform ‘uninformative’ priors on 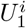 and 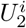 (note that these are so called ‘improper priors’). We also have that

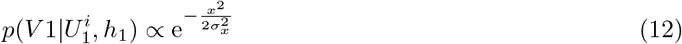

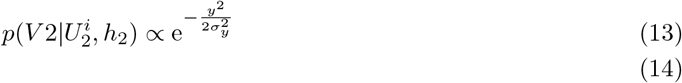

where

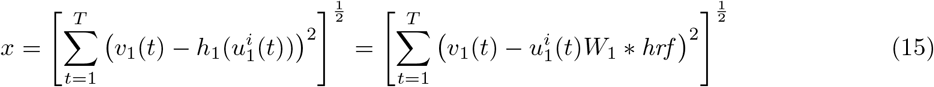

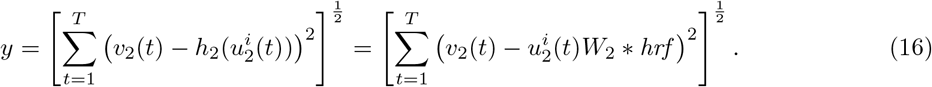

In the end

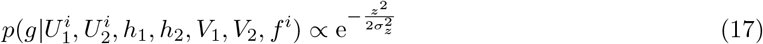

where

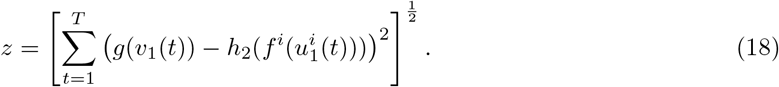

In conclusion,

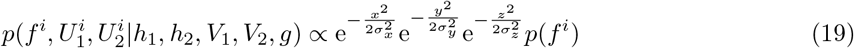

The prior *p(f^i^)* was taken as an uninformative prior, hyperparameter *σ_x_* was set to the total variation of *V*_1_, and hyperparameters *σ_y_* and *σ_z_* were set to the total variation of *V*_2_. The probability 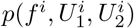 can be estimated marginalizing over the possible choices of *h*_1_ and *h*_2_. Two models of the underlying neural computations 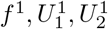 and 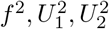 can be compared using the Bayes factor:

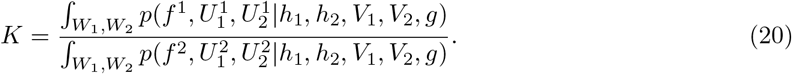

To simplify the comparison between models, *W_1_* and *W*_2_ were estimated as the parameters of a linear regression model with *U*_1_ * *hrf* and *U*_2_ * *hrf* as predictors (thanks to the associative property of convolution). To mitigate overfitting, *g* was estimated with all runs minus one, and then used to compute *z* in the independent, left-out run (following a leave-one-out procedure analogous to that used for the analysis of real fMRI data). This procedure was iterated for each choice of the left-out run. The probability of a model given the whole experiment was computed as the product of the probabilities for the individual runs. We tested 1) whether this approach could select the correct model with the responses in *C*_1_ as 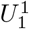 and the responses in *C*2 as 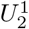, from an alternative model that also had the responses in *C*1 as 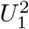, but had as 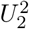 the responses in a second layer *C*2* of the same size as *C*2 constructed with linear transformations of the nodes in *C*1 (rather than the nonlinear pooling and max operation that generates *C*2); and 2) whether estimating a nonlinear *g* with artificial neural networks (nonlinear dependence) increased the probability assigned to the correct model 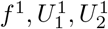 as compared to estimating *g* with multivariate linear regression (linear dependence).

### C. Experimental design

In the first experiment, eight participants (four female, mean age: 22 years, range: 19-25; all righthanded) completed three scanning sessions of ~ 2 hours each. Participants heard 165 two-second sounds frequently encountered in day-to-day life [14] and performed a sound intensity discrimination task. To download all 165 sounds, see the McDermott lab website:http://mcdermottlab.mit.edu/downloads.html. Sounds were presented using a block design, each block consisted of five presentations of an identical two-second sound, with one presentation 7 dB lower than the other four. All participants provided informed consent, and the Massachusetts Institute of Technology Committee on the Use of Humans as Experimental Subjects approved experiments. The fMRI data analyzed here is a subset of the data in [14], only including the subjects who completed three scanning sessions.

In the second experiment 11 participants completed a face localizer and a voice localizer. In the face localizer, participants watched 16s blocks of images of faces and houses while performing a 1-back task. In the voice localizer, participants performed a 1-back task on 16s blocks of voice and tool sounds. After the localizers, participants were administered five experimental runs in which they were asked to detect a target person identity, defined before the beginning of the experiment. Images of faces and recordings of voices were presented in 4s long trials arranged in a pseudorandomized order generated with Optseq 2 (http://surfer.nmr.mgh.harvard.edu/optseq/). Participants were asked to detect a famous target identity. Each run consisted of 120 trials and lasted approximately 8 minutes. All participants provided informed consent, and the IRB of Harvard University and the Ethics Committee of the University of Trento approved experiments.

In the third experiment, 16 participants completed a language localizer in which they passively read 18 seconds long blocks of sentences or nonwords. Each block consisted of 3 sequences of nonwords or of words forming a sentence. At the end of each sequence a cue was presented and participants had to press a button. After the localizer, participants listened to auditorily presented stories. Each run contained a full story, lasting approximately 6 minutes. In two of the runs, participants listened to the stories passively; in two other runs they performed a two-back task on the orientation of a line. All participants provided informed consent, and the Massachusetts Institute of Technology Committee on the Use of Humans as Experimental Subjects approved experiments.

### D. Data acquisition

Data for the first experiment were collected on a Siemens Trio 3T scanner with a 32-channel head coil at the Athinoula A. Martinos Imaging Center at McGovern Institute for Brain Research at MIT. Before collecting functional data, a high-resolution (1 × 1 × 1 *mm*^3^) T1-weighted MPRAGE sequence was performed (sagittal slice orientation, centric phase encoding, image matrix = 256 × 224 [Read × Phase], field of view =256 × 224 mm [Read × Phase], 128 axial slices, GRAPPA acquisition with acceleration factor = 2, duration = 5.36 minutes, repetition time = 2530, echo time = 3.48, TI = 1020 msec, 7 deg flip angle). Each functional volume consisted of fifteen slices oriented parallel to the superior temporal plane, covering the portion of the temporal lobe superior to and including the superior temporal sulcus. Repetition time (TR) was 3.4 seconds (acquisition time was only 1 second), echo time (TE) was 30 milliseconds, and flip angle was 90 degrees. For each run, the five initial volumes were discarded to allow homogenization of the magnetic field. In-plane resolution was 2.1 × 2.1 mm, and slice thickness was 4 mm with a 10% gap, yielding a voxel size of 2.1 × 2.1 × 4.4 mm. iPAT was used to minimize acquisition time. T1-weighted anatomical images were collected in each subject (1mm isotropic voxels) for alignment and surface reconstruction.

Data for the second experiment were collected on a Bruker BioSpin MedSpec 4T at the Center for Mind/Brain Sciences (CIMeC) of the University of Trento using a USA Instruments eight-channel phased-array head coil. Before collecting functional data, a high-resolution (1 × 1 × 1 mm^3^) T1-weighted MPRAGE sequence was performed (sagittal slice orientation, centric phase encoding, image matrix = 256 × 224 [Read × Phase], field of view =256 × 224 mm [Read × Phase], 176 partitions with 1mm thickness, GRAPPA acquisition with acceleration factor = 2, duration = 5.36 minutes, repetition time = 2700, echo time = 4.18, TI = 1020 msec, 7deg flip angle). Functional data were collected using an echo-planar 2D imaging sequence with phase oversampling (image matrix = 70 × 64, repetition time = 2000 msec, echo time = 21 msec, flip angle = 76 deg, slice thickness = 2 mm, gap = 0.30 mm, with × 3 mm in plane resolution). Over four runs, 1260 volumes of 43 slices were acquired in the axial plane aligned along the long axis of the temporal lobe.

Data for the third experiment were collected on a Siemens Trio 3T scanner with a 32-channel head coil at the Athinoula A. Martinos Imaging Center at McGovern Institute for Brain Research at MIT. Before collecting functional data, a high-resolution (1 × 1 × 1 *mm*^3^) T1-weighted MPRAGE sequence was performed (sagittal slice orientation, centric phase encoding, image matrix = 256 × 224 [Read × Phase], field of view =256 × 224 mm [Read × Phase], 128 axial slices, GRAPPA acquisition with acceleration factor = 2, duration = 5.36 minutes, repetition time = 2530, echo time = 3.48, TI = 1020 msec, 7 deg flip angle). Functional data were collected using an echo-planar 2D imaging sequence (image matrix = 96 × 96, repetition time = 2000 msec, echo time = 30 msec, flip angle = 90 deg, 31 slices, slice thickness = 4mm, 10% distance factor, with 2.1 × 2.1 mm in plane resolution). Prospective acquisition correction was used to adjust the positions of the gradients based on the participant’s motion one TR back. The first 10s of each run were excluded to allow for steady-state magnetization.

### E. Data analysis

#### 1. Preprocessing

The data of the first experiment were preprocessed using FSL and in-house MATLAB scripts. Volumes were corrected for motion and slice time. Volumes were skull-stripped, and voxel time courses were detrended linearly. Alignment of functional volumes to the anatomical volume was performed with FLIRT [15] and BBRegister [16]. The resulting functional volumes were then resampled to vertices on the reconstructed cortical surface computed via FreeSurfer [17, 18], and were smoothed on the surface with a 3mm FWHM 2D Gaussian kernel to improve SNR. Analyses for the first experiment were done in this surface space, but for ease of discussion we refer to vertices as ‘voxels’ throughout this paper. For each scan sessions, we estimated the mean response of each voxel (in the surface space) to each stimulus block by averaging the response of the second through the fifth acquisitions after the onset of each block (the first acquisition was excluded to account for the hemodynamic lag). Pilot analyses showed similar response estimates from a more traditional GLM. Responses were converted to percent signal change (PSC) by subtracting and dividing by each voxel’s response to the blocks of silence, and downsampled from the surface space to a 2mm isotropic grid on the FreeSurfer-flattened cortical sheet. For individual subject analyses, we used the same voxel selection criterion as [14] selecting voxels with a consistent response to sounds from a large anatomical constraint region encompassing the superior temporal and posterior parietal cortex.

The data of the second and third experiments were preprocessed with SPM12 (http://www.fil.ion.ucl.ac.uk/spm/software/spm12/) running on MATLAB 2015b. After slice-timing correction and realignment, the functional volumes were coregistered to the anatomical volume and normalized. No smoothing was applied. Functional regions of interest (ROIs) were defined in individual subjects with t-contrasts in the functional localizers for faces¿houses and voices¿tool sounds in the first experiment, for sentences¿nonwords in the second experiment. In the first experiment, ROIs were defined as 9mm radius spheres centered in the t-contrast peaks in the anatomical areas where activations were expected based on prior studies. In the second experiment, ROIs were defined by taking the 100 voxels showing highest t-contrasts within each of six search spaces generated based on a probabilistic activation overlap map for the localizer contrast in 220 participants [14]; the two anterior temporal and two posterior temporal search spaces were morphed together. Patterns of response in each ROI were extracted and denoised by scrubbing outliers with ART (RRID:SCR_005994) and then applying CompCor [19], regressing out the first five principal components extracted from a white matter and cerebrospinal fluid mask defined in individual participants.

#### 2. NL-MVPD

For each region of interest, the mean timecourse was subtracted from each voxel, therefore removing the mean univariate response in each region at each timepoint. In the first experiment, all runs were analyzed together, in the second experiment, the runs with passive listening and the runs with the active task were analyzed separately. Independent sets of data were generated following a leave-one-run-out approach. The training data were used to perform dimensionality reduction using PCA, where the number of principal components was determined using the Bayesian Information Criterion (BIC). The testing data were then projected on the dimensions estimated using the training data. This procedure was identical for the linear and for the nonlinear models. For the linear models, the mappings between each pair of brain regions were modeled with a multiple regression taking as inputs the values along the dimensions in one brain region in the pair and taking as outputs the values along the dimensions in the other region. For the nonlinear models, mappings were modeled with one-hidden-layer artificial neural networks with the same inputs and outputs. The networks used all-to-all connectivity between layers, hyperbolic tangent transfer functions and were trained with backpropagation using the Levenberg-Marquardt algorithm.

All models were estimated using the training data, and variance explained was calculated in the left-out run that was not used for training as the square of the correlation between the predicted and observed timecourses for each component, subsequently averaging across the components. Note that calculating the correlation between a regions response and its nonlinear prediction based on another region’s response is closely related to mutual information, but thanks to this framework, unlike with mutual information, we can generate predictions for independent data. This procedure (PCA and model training/testing) was repeated for all choices of the left-out run. In order to facilitate comparison with functional connectivity measures, which use *r* values rather than R2, the squared root of variance explained was computed for each dimension and averaged, obtaining a ‘generalized correlation’ index |*r*|. Thanks to the independence of the training/testing procedure, the estimates of |*r*| are not biased by the complexity of the models, and in fact on average they asymptote for a small number of hidden nodes (Supplementary Figures 4, 7). Network visualization on the cortex was performed with BrainNet Viewer [20]. The nonlinear shift was calculated applying the Fisher transformation to the |*r*| matrices for linear and nonlinear MVPD (to normalize the data), and subtracting the two Fisher-transformed matrices. Adelson-Darling tests did not detect significant deviation from normality of the distribution of nonlinear-shift (identity: *p* = 0.3484; language, no cognitive load: *p* = 0.3467; language, cognitive load: *p* = 0.4528). Nonmetric multidimensional scaling was performed with the MATLAB function ‘mdscale’ using default parameters. Comparison between nonlinear integration between modalities and nonlinear integration within modality only (Fig. 2F) was performed subtracting the maximum |*r*| values for each of the model types within a range between 1 and 20 hidden nodes and assessing the significance of the difference across participants with a Wilcoxon signed-rank test. In the language experiment, classification of task based on connectivity matrices was performed with linear discriminant analysis (LDA) using a leave-one-subject-out cross validation procedure. Statistical significance was assessed with a permutation test using 1000 iterations.

**Figure 2:**
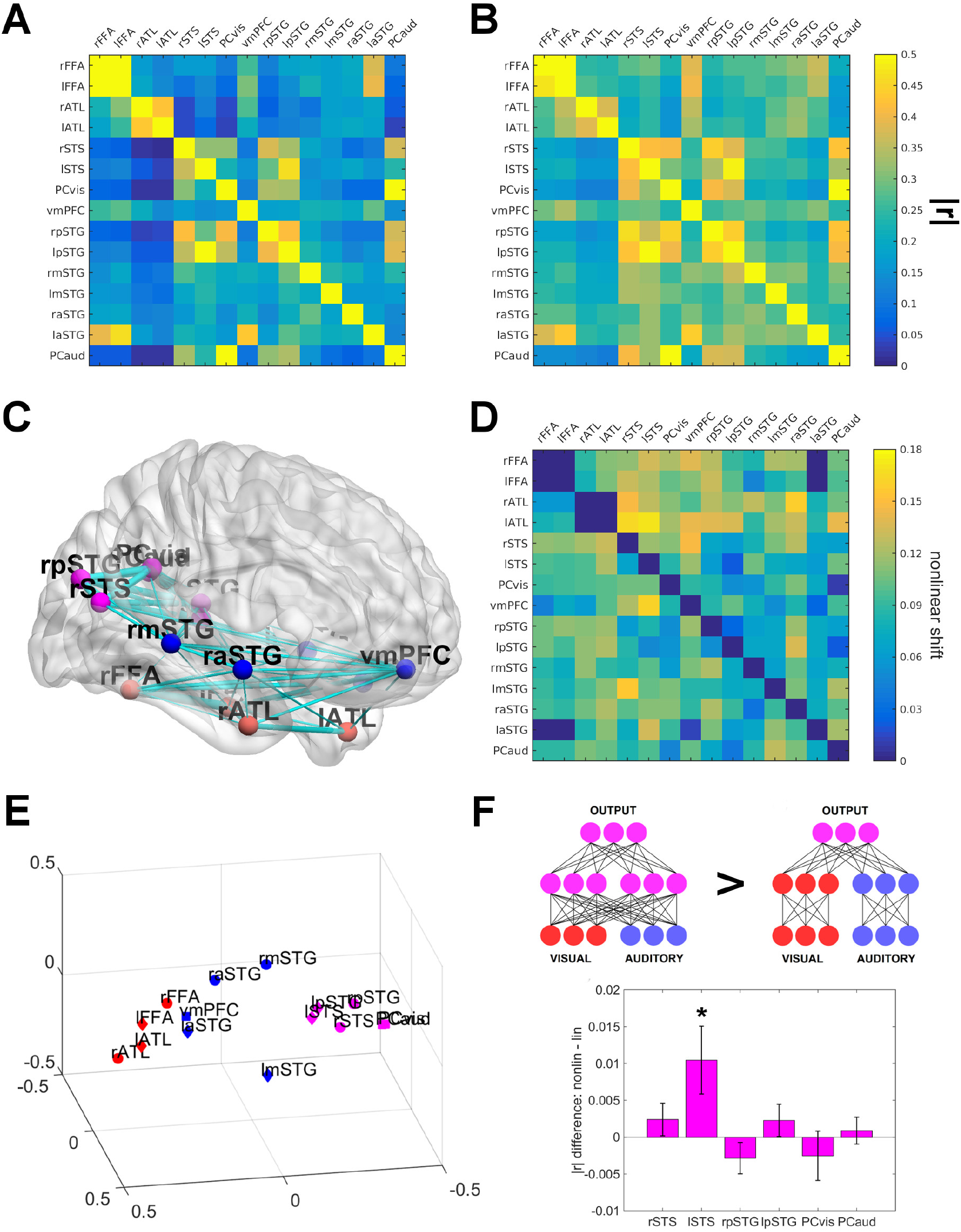
Linear and nonlinear pattern dependence between face- and voice-selective regions. A) Linear multivariate dependence matrix generated with multiple regression. B) Nonlinear multivariate dependence matrix generated using the same data with a one-hidden-layer artificial neural network with 5 hidden nodes. C) Visualization of the structure of nonlinear dependence on the cortical surface. D) Nonlinear-shift: increase in statistical dependence produced by nonlinear vs linear models for each pair of regions. E) Multidimensional scaling based on nonlinear multivariate dependence matrix shows clustering of face-selective regions (red), voice-selective regions (blue) and regions with overlap between face- and voice-selectivity (purple). F) Example of nonlinear dependence for one individual subject between one pair of regions, restricted to the first 2 PCs of the predictor region (FFA) and the first PC of the target region (vmPFC) to be visualized in 3 dimensions. The surface depicts the nonlinear dependence estimated using the training data (blue dots), it folds downward on the right side better predicting independent testing data (red dots). Better performance of the nonlinear model is reliable across participants (see panel D).

## III. RESULTS

As a first test, we asked whether NL-MVPD can select the correct underlying model of nonlinear neural computations, when the ground truth is known. To do so, we simulated fMRI data drawn from two layers of a widely-used neural network model of object recognition (HMAX, [3]). The responses in the C1 and C2 layers of HMAX were calculated for 10 standard images. FMRI responses were generated by (i) considering each voxel as a linear combination of randomly sampled nodes, (ii) convolving with a haemodynamic response function, and (iii) adding gaussian noise (see Supplementary Materials). One region (200 voxels) was generated from C1 nodes and a second region from C2 nodes. We then tested whether NL-MVPD could correctly select the underlying nonlinear dependence, compared to an alternative hypothesis of linear dependence between C1 and C2 nodes (Supp Fig. 1). NL-MVPD correctly recovered the nonlinear dependence in the underlying neural computations from the simulated fMRI voxel responses (Bayes factor *K* > 20) and assigned greater probability to the true model than linear dependence (Bayes factor *K* > 20). Second, we asked whether NL-MVPD provides a better fit to real fMRI data drawn from regions that are very likely to be related via non-linear multivariate statistical dependence. Participants (n=8) listened to 165 clips of natural sounds, including environmental sounds (toilet flushing, rain), music, and human speech. We estimated the multivariate timecourses in two regions of auditory cortex (Supp. Fig. 2A). These regions were identified in a previous stationary analysis, as having responses better explained by an earlier or a later layer in a CNN, and they corresponded, respectively, with portions of primary and non-primary auditory cortex. Here we asked whether the dependence between these regions was better captured by NL-MVPD than by standard measures of linear or univariate functional connectivity. As predicted, the dependence between these regions of auditory cortex was better captured by NL-MVPD than when using alternative models of inter-regional dependence. In this case, neural networks with relatively few hidden nodes (1 to 5) performed better than networks with more nodes (6 to 10), showing that exceedingly complex models can lead to overfitting (Supp. Fig. 2B). Third, we tested whether NL-MVPD could be used to capture inter-regional dependences within a larger and more diverse network of regions involved in face- and voice-identity recognition. Participants (n=11, 6 female) viewed the faces and listened to the voices of three highly-recognizable and famous national politicians. Previous stationary analyses (16) identified cortical regions that contain multivariate information about face identity, voice identity, and person-identity generalizing across stimulus modality (Supp. Fig. 3).

Here we measured the statistical dependence between these regions (Fig. 2A-C). Overall, NL-MVPD (Fig. 2A better captured the variance in left-out data than a model of only linear dependence between regions (Fig. 2B) or a model of task-driven activity alone (Supplementary Figure 4); the fit peaked for neural networks with 4 hidden nodes. We measured the increase in statistical dependence produced by nonlinear, versus linear, models for each pair of regions (‘nonlinear shift’). The largest nonlinear shift was between regions implicated in encoding increasingly invariant representations of identity (posterior STS - anterior temporal lobe, Fig. 2D, 16). By contrast, almost no nonlinear shift was observed for pairs of homologous regions in opposite hemispheres (e.g. right and left FFA, right and left ATL), consistent with the expectation that homologous regions play similar functional roles, with no intervening nonlinear computations. We then sought to characterize the network structure for person-identity processing revealed by NL-MVPD. Consistent with expectations, we identified stronger statistical dependence between pairs of brain regions responsive to the same modality (i.e., among face-processing regions, and among voice-processing regions, Fig. 2E). Multi-dimensional scaling (MDS) was used to visualize the structure of the functional network (using closeness to represent the strength of statistical dependence, Fig. 2F). Homologous regions in the two hemispheres were arranged close to each other in the MDS space, and in similar relative positions with respect to the other regions in their hemisphere, indicating a common functional organization across the two hemispheres. Responses in a set of brain regions showing multimodal responses (visual and auditory) were modeled as a joint nonlinear function of the responses in unimodal visual regions and unimodal auditory regions. We compared a model in which nonlinearities only occur within the separate modalities, to a model with nonlinear integration of the two modalities (Fig. 2F). Nonlinear integration between different modalities generated better predictions in the left posterior superior temporal sulcus (lSTS, Wilcoxon Signed Rank test: *p* = 0.0322). Fourth, we tested the application of NL-MVPD to complex language comprehension, for which animal models of inter-regional dependence are unlikely, making fMRI data pivotal. Language comprehension is known to engage a network of frontal and temporal brain regions [21], but the dependence between regions during language comprehension has been difficult to resolve. Participants listened to four verbal narratives; two while passively listening, and two while simultaneously performing an unrelated cognitively demanding task (a two-back task on the orientation of a line). Using an independent dataset [22], language selective regions were defined in individual participants (Supp. Fig. 6), bilaterally in the frontal lobe (IFGorb, IFG, and MFG), and in the temporal and parietal cortices (AntTemp, PostTemp, and AngG). NL-MVPD again provided a better predictive model of statistical dependence between brain regions than linear models, for brain activity measured during both passive listening and listening under cognitive load (Supplementary Figure 7).

We then sought to characterize the network structure for language comprehension revealed by NL-MVPD. The statistical dependence matrix (Fig. 3A-B) and multi-dimensional scaling (Figure 4A) during the passive listening task suggested four candidate subnetworks: 1) bilateral posterior temporal - MFG, 2) bilateral IFGorb, 3) bilateral IFG, and 4) bilateral AngG - AntTemp. In three of these subnetworks (1,3, and 4), |*r*| values for pairs of regions within a subnetwork were significantly greater than |*r*| values for pairs of regions belonging to different subnetworks across participants (Wilcoxon Signed Rank test for the four subnetworks: *p* = 0.0013, p = 0.1788, *p* = 0.0005, *p* = 0.0011; Fig. 4B). Furthermore, all four candidate subnetworks identified in the passive listening data were replicated in the cognitive load data (Wilcoxon Signed Rank *p* = 0.0494, *p* = 0.0032, *p* = 0.0019, *p* = 0.0027; Fig. 4C-D).

**Figure 3:**
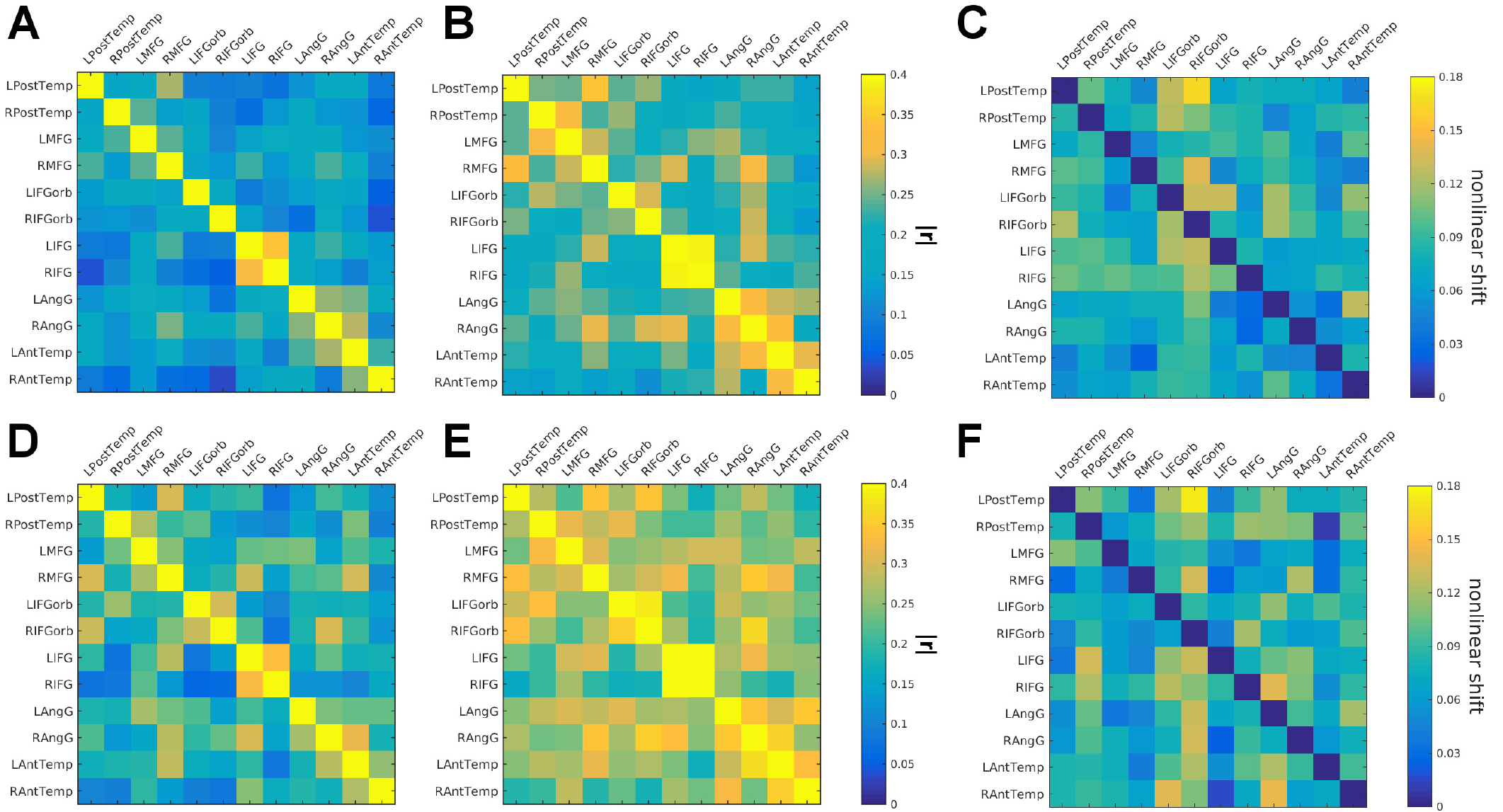
Linear and nonlinear multivariate pattern dependence between language-selective regions. A) Linear multivariate dependence matrix in the passive listening task. B) Nonlinear multivariate dependence matrix for 3 hidden nodes in the passive listening task. C) Nonlinear-shift: increase in statistical dependence produced by nonlinear vs linear models for each pair of regions in the passive listening task. D) Linear multivariate dependence matrix in the cognitive load task. E) Nonlinear multivariate dependence matrix for 3 hidden nodes in the cognitive load task. F) Nonlinear-shift: increase in statistical dependence produced by nonlinear vs linear models for each pair of regions in the cognitive load task.

**Figure 4:**
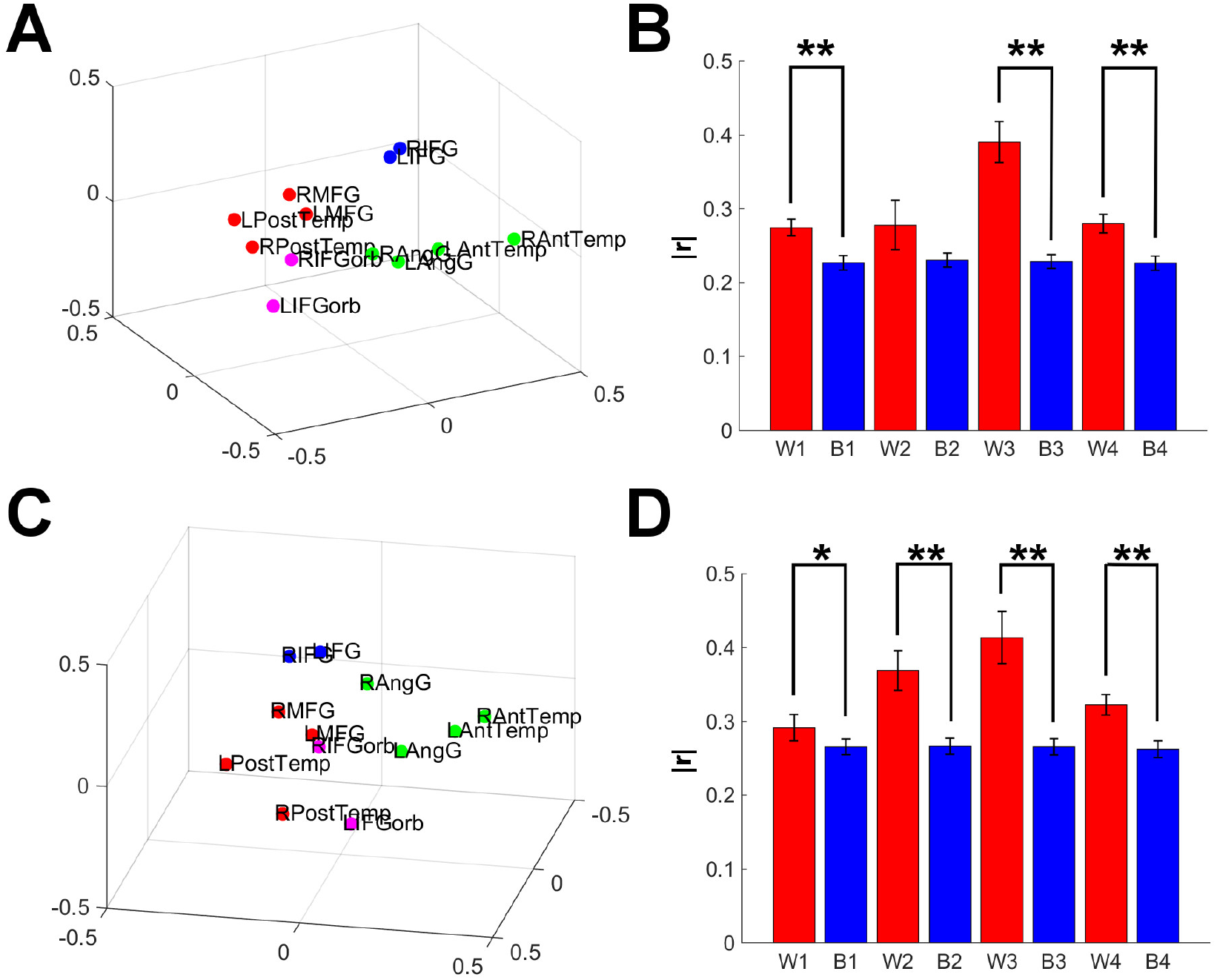
Multidimensional scaling and subdivision of the language network into subnetworks that are reliable across participants. A) 3D multidimensional scaling based on the nonlinear dependence matrix of the passive listening task. B) Average nonlinear dependence —r— values for pairs of regions within (W) and between (B) the subnetworks in the passive listening task: PostTemp-MFG (’W1’, ’B1’), IFGorb (’W2’,’B2’), IFG (’W3’,’B3’) and AngG-AntTemp (’W4’,’B4’). C) 3D multidimensional scaling based on the nonlinear dependence matrix of the two-back task. The brain regions are arranged similarly in space across the two tasks. D) Average nonlinear dependence —r— values for pairs of regions within and between the subnetworks in the cognitive load task: PostTemp-MFG (’W1’, ’B1’), IFGorb (’W2’,’B2’), IFG (’W3’,’B3’) and AngG-AntTemp (’W4’,’B4’).

To the extent that statistical dependence between brain regions reflects the underlying neural processing, it should be different when participants are engaged in different tasks. To test whether the difference between passive listening and listening under cognitive load was reflected in nonlinear dependence, linear discriminant analysis (LDA) was applied to the nonlinear mappings between regions in the two tasks, using as inputs the corresponding statistical dependence matrices. An LDA classifier was trained with data from all but one participant, and the accuracy at classifying the tasks in the left-out participant was assessed. This procedure yielded a mean classification accuracy of 62.5% (*p* = 0.031, permutation test, 1000 iterations). Linear dependence did not yield significant classification (accuracy = 56.25%, *p* = 0.144, permutation test, 1000 iterations), nor did standard functional connectivity (accuracy = 43.75%, *p* = 0.739, permutation test, 1000 iterations). During the cognitive load task, the nonlinear dependence between subnetworks increased as compared to the nonlinear dependence within subnetworks (*t*(15) = 2.65, *p* = 0.0182), indicating greater intercommunication between the subnetworks under cognitive load.

## IV. DISCUSSION

Nonlinear multivariate pattern dependence (NL-MVPD) between brain regions can be measured in humans, in typical fMRI datasets. In simulated data and in three experiments with different stimuli and tasks, nonlinear multivariate models of dependence outperformed linear models in predictions for left-out data. In the second experiment, nonlinear multivariate dependence identified expected modality-dependent structure in the network of face- and voice-selective regions. In the third experiment, nonlinear multivariate dependence identified a subdivision of the language network into distinct and reliable subnetworks, and revealed modulations in the interactions between brain regions driven by the task performed by participants.

Being sensitive to interactions between brain regions that could not be captured with standard univariate or linear methods is not the only advantage of NL-MVPD. Using imaging data to constrain computational theories of cognition has been a longstanding challenge in Neuroscience, with critics highlighting the limitations of the inferences about cognition derived from fMRI experiments [23]. Insights from functional imaging have inspired theoretical work [24] on object recognition, and neural patterns of responses have been used to constrain theories of the properties encoded by a brain region [25], but a systematic approach to model neural computations based on fMRI data has yet to be developed. By capturing the multivariate and nonlinear dependence between brain regions, NL-MVPD brings us closer to the goal of using fMRI evidence to constrain computational models of cognition. To outline a possible path towards that goal, the approach used in the simulation-based validation in this article could be extended to perform model search on a larger space of neural networks, thus using the fMRI data to generate high-probability neural network models.

Since NL-MVPD predicts responses in a brain region based on responses in other regions, it can explain reliable variance in neural responses which is not driven by the concurrent external stimuli, but which is instead driven by internal states like endogenous attention shifts, or spontaneously retrieved memories. As a consequence, NL-MVPD has the potential to complement forward models of brain activity which rely entirely on stimulus properties [26].

NL-MVPD can be applied even to tasks for which models of the underlying neural computations (such as Deep Neural Networks) are not available. These properties make NL-MVPD very flexible and complementary to existing analyses techniques.

NL-MVPD stands at the confluence between existing techniques for multivariate pattern analysis (MVPA) and for measuring functional connectivity. Introducing nonlinearity in the statistical dependence, NL-MVPD differs from previous approaches to integrate MVPA and connectivity [8–12]. An additional difference between NL-MVPD and informational connectivity [9] is that NL-MVPD characterizes each region’s responses in a data-driven fashion to maximally capture their variance, while informational connectivity is based on experimenter-defined classifications. The two approaches are therefore complementary, each offering distinct advantages to address different questions. As compared to other work that integrates MVPA and connectivity [8–12], the method described in this article leverages the potential offered by Multivariate Pattern Connectivity (MVPC, [11]) in which the maps between brain regions are themselves multivariate, and combines it with nonlinear functions in order to predict the responses in one region as a function of multiple dimensions represented in another region. For example, one region might encode view-specific information about face parts (e.g. different dimensions encode the shape, size or distance of one feature from the others); another region might encode view-independent information about face identity; the proposed method has the potential to measure the nonlinear transformation of dimensions in the space of parts required to predict responses in the space of face identity.

The results show that this approach is not only theoretically principled, but also practically viable. Non-linear mappings between brain regions explain more variance in independent data than linear mappings, demonstrating that even the limited data obtained with a standard fMRI protocol justify the use of nonlinear models. In other words, the data are sufficiently rich that nonlinearity can be introduced without leading to overfitting or compromising generalization to independent data.

It is important to note that variance explained by a nonlinear mapping between two regions cannot be used to infer the presence of a direct anatomical connection between the two regions. In fact, like in other methods based on statistical dependencies between functional responses (e.g. functional connectivity), high |*r*| values can potentially be observed even when the interaction between two regions is mediated by a third region.

Several lines of research can be pursued in the future to increase the potential of this modeling approach. A critical step will be to relate the principal components of responses in each individual region to properties of the stimuli (i.e. an encoding model of each region’s representational space [26]). Next, data driven models of nonlinear mappings between brain regions could be used to constrain algorithmic models of neural computation. In the end, graph analysis methods [27] could be used to study the structure of networks composed of nonlinear mappings. From a computational perspective, the goal of connectivity research includes understanding how information is transformed from brain region to brain region. By capturing properties of neural computations like their multivariate and nonlinear nature, NL-MVPD marks a key step towards this goal. NL-MVPD is generally applicable to a wide variety of domains, including the study of clinical populations. We hope that nonlinear statistical dependence will help to better understand and potentially diagnose neurological disorders. In the end, NL-MVPD is not limited to fMRI, but can be applied to data obtained with other techniques, like multi-site electrophysiological recordings in animals.

## Acknowledgments

This work was funded by NIH Grant 1R01 MH096914-01A1 to Prof. Rebecca Saxe, by a grant from the NICHD to EF (HD057522), a grant from the Simons Foundation to the Simons Center for the Social Brain at MIT to E.F. and R.S., and by the Center for Mind/Brain Sciences at the University of Trento. We also thank the support of a fellowship from the Simons Center for the Social Brain to S.A., and of a DOE Computational Science Graduate Fellowship (DE-FG02-97ER25308) to A.K. We would like to thank Josh McDermott and Sam Norman-Haignere for making available the auditory data; Josh Tenenbaum for comments on an earlier draft of the manuscript, and Tyler Bonnen, Dae Houlihan, Zach Mineroff, Alex Paunov, Valentina Brentari and Briana Pritchett for assistance during data acquisition and implementation of experiment scripts. We would like to acknowledge the Athinoula A. Martinos Imaging Center at McGovern Institute for Brain Research at MIT, and the support team (Steve Shannon, Atsushi Takahashi, and Sheeba Arnold). The data are available from the corresponding author upon request.

